# Protoxylem microtubule patterning requires ROP pattern co-alignment, realistic microtubule-based nucleation, and moderate microtubule flexibility

**DOI:** 10.1101/2024.04.04.588070

**Authors:** Bas Jacobs, Marco Saltini, Jaap Molenaar, Laura Filion, Eva E. Deinum

## Abstract

The development of the water transporting xylem tissue in plants involves an intricate interplay of Rho-of-Plants (ROP) proteins and cortical microtubules to generate highly functional secondary cell wall patterns, such as the ringed or spiral patterns in early-developing protoxylem. We study the requirements of protoxylem microtubule band formation with simulations in CorticalSim, extended to include finite microtubule persistence length and a novel algorithm for microtubule-based nucleation. We find that microtubule flexibility is required to facilitate pattern formation for all realistic degrees of mismatch between array and pattern orientation. At the same time, flexibility leads to more density loss, both from collisions and the microtubule-hostile gap regions, making it harder to maintain microtubule bands. Microtubule-dependent nucleation helps to counteract this effect by gradually shifting nucleation from the gap regions to the bands as microtubules disappear from the gaps. Our results reveal the main mechanisms required for efficient protoxylem band formation.

## 1 Background

To meet the water demand of their above ground organs, plants depend on their water transporting tissue, the xylem. Xylem consists of an interconnected tubular network that stretches from young root tips to the above ground water sinks. Inside the xylem, the water pressure is often negative [1], meaning that the constituting elements have to be sufficiently strong to prevent collapse [2]. This is extra challenging close to the root tips. The earliest maturing xylem vessels, called protoxylem, mature while the surrounding tissue is still elongating [3]. Vessel elements could not stretch along if they have homogeneously thick cell walls. Therefore, protoxylem elements show banded or spiralled secondary cell wall reinforcements. These bands provide strength [4], but also allow for elongation. In contrast, the later maturing metaxylem has much more solid secondary cell wall reinforcements, with a number of ellipsoid gaps [5]. More general, these two xylem types are studied as a model system for pattern formation in cell wall structure [6].

The patterned secondary cell wall reinforcements initially consist mostly of cellulose microfibrils deposited by cellulose synthase (CESA) complexes [5, 7]. Upon maturation, they are additionally lignified [8]. CESA delivery to the membrane and their subsequent movement through it is guided by the cortical microtubule array [9–13], with the cellulose microfibrils in the cell wall themselves as a secondary directing mechanism in absence of microtubule contact [13]. Understanding how the characteristic protoxylem cell wall patterns form, therefore, requires understanding how the corresponding patterns arise in the cortical microtubule array.

Microtubule dynamics during xylem patterning can be studied by ectopic expression of the transcriptional master regulators VASCULAR NAC DOMAIN (VND) 6 and 7, which induce metaxylem and protoxylem like patterns, respectively [14, 15]. In these systems, the cortical microtubule array adopts the pattern that the subsequent secondary cell wall reinforcements will follow [16, 17]. This pattern is formed in interaction with Rho-of-Plants (ROP) proteins and their downstream effectors. In metaxylem, active AtROP11 accumulates in future gap regions, where it recruits MICROTUBULE DEPLETION DOMAIN 1 (MIDD1) and Kinesin-13A, leading to local microtubule depolymerization [18–20]. In protoxylem, a similar interaction between ROPs and microtubules is highly likely, and striated ROP and MIDD1 patterns have been observed [17, 21]. Many ROPs are expressed in the zone of protoxylem patterning [22], however, and the ROPs responsible for the protoxylem pattern remain elusive as even *AtROP7/8/11* triple knockouts still form a banded protoxylem pattern [17], possibly due to a high degree of redundancy. Microtubules themselves also influence the shape of the ROP pattern by anisotropically restricting active ROP diffusion [19, 23], which can orient the ROP pattern along the microtubules [24]. This coupling results in a degree of co-alignment between the orientation of the original microtubule array and the banded ROP pattern, particularly in highly aligned arrays. Previous simulations suggest that this co-alignment increases the speed of pattern formation [16], but it has not been thoroughly quantified to what degree of co-alignment is required. Consequently, we do not know the likelihood that individual arrays meet this requirement in practice.

Individual microtubules are highly dynamic, particularly at their plus-end (figure 1A). They display phases of growth and rapid shrinkage, with stochastic switches between the two, called catastrophe and rescue [25–27]. As the cortical microtubules are attached to the inside of the cell membrane, they are bound to interact via frequent collisions. The outcome of these collisions depends on the relative angle of the colliding and obstructing microtubule [28]. For small angles, the colliding microtubule bundles with the obstructing one, while for large angles it either crosses over or undergoes and induced catastrophe [28]. After a crossover (figure 1B), the latest arriving microtubule, i.e., on the cytoplasmic side, is most likely to be severed by katanin at a later time (figure 1B) [29, 30]. Computer simulations and theoretical models have been indispensable in understanding how the above interactions can lead to the spontaneous self-organization of the cortical array into highly aligned patterns [31–39]. As computer simulations are more easily amended to complex situations, they have been the approach of choice in studying protoxylem development [16, 40]. Full reproduction of the patterning process with interacting microtubules, however, turned out to be a hard problem and has not been achieved yet [16].

**Fig. 1.**
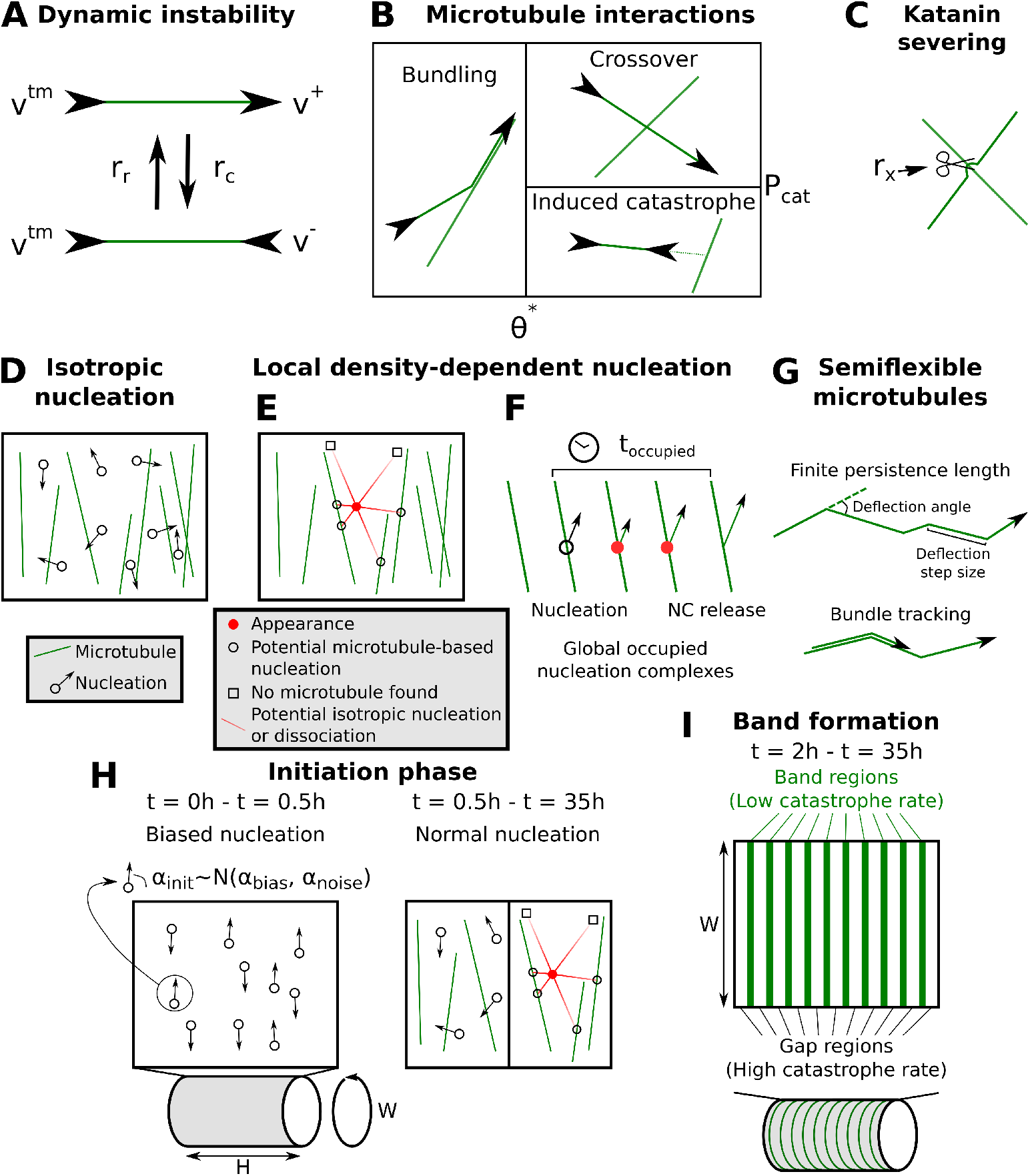
Microtubule simulation details. (A) Microtubule plus ends grow or shrink at constant rates *v*^+^ or *v*^*−*^, respectively, and minus ends retract at constant rate *v*^*tm*^. Spontaneous catastrophes (switch from growing to shrinking) occur at a rate *rc* and rescues (switch from shrinking to growing) at a rate *rr* . (B) Microtubule-microtubule collision outcomes depend on the collision angle. At angles below *θ*^*∗*^ the impinging microtubule bundles with, i.e., continues growing along, the obstructing one. At greater collision angles, the incoming microtubule undergoes an induced catastrophe with a probability *Pcat* and crosses over the other microtubule otherwise [31, 52]. (C) Katanin severs the overlying microtubule at crossovers at a constant rate *rx* per crossover. (D) With isotropic nucleation, new microtubules appear at a constant rate *rn*, in a uniformly random location and direction. (E) With local density-dependent nucleation, new microtubules appear in a way that more realistically mimics the behaviour of nucleation complexes (NCs), taking into account their appearance at the membrane, diffusive movement and potential microtubule binding. (F) In addition, nucleation complexes stay occupied for a duration *t*_*occupied*_, temporarily reducing the global nucleation rate. (G) For simulations with semiflexible microtubules, a finite persistence length is achieved via discrete deflections in the microtubule growth direction. In addition, microtubules in bundles follow their bundle around bends below an angle *θ*_*b*_ (‘bundle tracking’). Deflection angles in cartoons are exaggerated for visibility. (H) In band formation simulations, an initial transverse array is artificially enforced by drawing nucleation angles in the first half hour of simulated time (*α*_*init*_) from a normal distribution with an average of *α*_*bias*_ and a variance of *α*_*noise*_. (I) Protoxylem band formation is simulated with predefined band and gap regions, where the catastrophe rate in the gap regions is increased by a factor *fcat* after a 2*h* initiation phase, following [16], except that *fcat* is reduced to 3.

One major reason is related to microtubule nucleation. Most microtubules in the cortical array are nucleated from existing microtubules with a specific distribution of relative nucleation angles [41]. The most common implementation of microtubule based nucleation [16, 32, 34], however, introduces a global competition for nucleation, leading to highly inhomogeneous arrays [40] and aggregation of microtubule density in one or few bands of the protoxylem pattern only [16, 40]. Recently, it has been shown that this so-called “inhomogeneity problem” is a consequence of an incomplete understanding of the nucleation process [40]. A more realistic nucleation algorithm based on detailed experimental observations is indeed sufficient to produce regular banded protoxylem arrays in a simplified context of transverse, non-interacting microtubules [40]. The question is whether the adoption of a more realistic nucleation algorithm [42] alone will be sufficient to produce timely band formation in the full interacting microtubule array.

Another potentially important aspect of microtubules that is often disregarded in modelling studies is their flexibility (figure 1G). Most cortical array studies model microtubule segments in between collision points as perfectly straight line [16, 31, 33, 36, 43, 44], justified by a millimetre range persistence length of isolated microtubules [45, 46]. In microscopic images of plant cells, however, microtubules appear less straight, e.g. [16, 47–49], suggesting a lower effective persistence length in the cell context. Unfortunately, the actual persistence length in plant cells has never been quantified yet, and the recent studies that do model microtubules as semiflexible polymers [37, 39], base themselves on measurements from animal cells [50, 51]. The values they use are similar to the values found *in vitro* for microtubules with reduced persistence length in the presence of MAP70-5, addition of which introduces small circular microtubule bundles [46]. Actual persistence length is, therefore, likely higher in transverse arrays and protoxylem bands. The effect of different degrees of flexibility on protoxylem patterning remains an open question. Flexibility may both help and hinder the patterning process, as incorrectly oriented microtubules may find their way back to a band, while correctly oriented ones may curve out of it.

Here, we study the effect of three different concepts on protoxylem microtubule patterning: (1) a co-alignment between microtubules and the microtubule-hostile future band regions likely specified by ROPs, (2) realistic microtubule-based nucleation through a recently developed computationally efficient nucleation algorithm that captures the critical aspect of localized positive feedback [42], and (3) a realistic degree of microtubule flexibility.

## 2 Methods

### 2.1 Microtubule simulations

We performed our simulations using an extended version of the cortical microtubule simulation software ‘CorticalSim’ [52], fast, event-driven software for simulating cortical microtubule dynamics and interactions on the cell surface (cortex) [16, 31, 34, 36, 52, 53]. Unless stated otherwise, we used a cylindrical geometry with dimensions representative of a developing protoxylem cell (a height of 60 *µm* and a radius of 7.5 *µm*). For a detailed overview of all parameter values, see Appendix A.

### 2.2 Microtubule dynamics

The microtubules are modelled as connected series of line segments that together have one plus end and a minus end. The minus end retracts at a constant speed *v*^*tm*^ (Fig 1A) simplifying observed minus end dynamics [54], while the plus end mimics dynamic instability by switching between growing and shrinking states (catastrophes and rescues, Fig. 1A). When a microtubule collides with another microtubule, this results in a bundling event for collision angles *θ < θ*^*∗*^ after which the microtubule continues growing along the obstructing microtubule, and an induced catastrophe or crossover event for larger angles (Fig. 1B). Additionally, when a microtubule collides with the edge of the bounding cylinder, it also undergoes a catastrophe, which helps favour a transverse array orientation [55]. Finally, any microtubule crossing over another can undergo a severing event at the intersection with a rate *r*_*x*_ per crossover, creating a new shrinking plus end and retracting minus end (Fig. 1C).

### 2.3 Microtubule flexibility

In order to model the underlying flexibility of the microtubules, we extended the CorticalSim model drawing inspiration from the method in [37]. Specifically, we introduced “deflection” events that abruptly change the microtubule growth direction. The deflection step size, i.e. the length a microtubule grows before the next deflection occurs, is drawn from an exponential distribution with mean *l* . The deflection angle is drawn uniformly from [*−m, m*], where *m* is the maximum deflection angle calculated to obtain the desired persistence length given 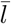 . Note that deflection angles with absolute values lower than a minimal deflection angle *q* = 0.1° were set to zero to avoid numerical artefacts. With these parameters we can control the persistence length *p* as shown in Appendix C. To prevent microtubules from leaving bundles at every deflection point, we made microtubules follow their bundle along bends smaller than 10° (Fig. 1G). For further implementation details, see Appendix B.

In our simulations, microtubule bundles consist of multiple microtubules on the same trajectory, without any space in between. Therefore, there is no distinction between microtubules on the edges of the bundle, which could in principle deflect outwards, and microtubules on the inside, that do not have any room to deflect at all. To approximately overcome this limitation, we reject a fraction *n/*(*n* + 1) of deflections in bundles, where *n* is the number of other microtubules in the bundle at the point of the deflection. Additionally, in bundles, we would expect most microtubules to stay with the bundles through small bends. To incorporate this feature, we force microtubules to track their bundles along bends, as long as the bending angle is not too large. Specifically, since bundling events create bend points and these may have large angles (up to 40°), we implemented a maximum bundle tracking angle of *θ*_*b*_ = 10°. If a bundle splits with an angle below this value, an incoming microtubule randomly follows one of the bundles, proportional to their occupancy.

### 2.4 Microtubule nucleation

In many microtubule modelling studies [31, 37–39] new microtubules appear by isotropic nucleation, i.e, with random, uniformly distributed locations and orientations (Fig. 1D). However, it has previously been shown that in practice, most microtubules are nucleated from existing microtubules, with a specific distribution of angles [41]. In our model, we explicitly include this nucleation mechanism, modelling the angle distribution following the approach of Deinum et al. [34]. Additionally, we perform some simulations with purely isotropic nucleation for comparison.

To avoid the inhomogeneity problem of previous microtubule-bound nucleation algorithms [like 34], we use a recently developed more realistic nucleation algorithm dubbed “local density-dependent nucleation” [42] that effectively approximates the diffusion of nucleation complexes from their appearance at the plasma membrane to the point where they either dissociate or nucleate. In this approach, nucleation complexes are not modelled as explicitly diffusing particles with additional dynamics, but handled implicitly via instantaneous appearance events, occurring at a rate *r*_ins_. During such an event, the algorithm draws an “appearance point” for a complex at a random, uniformly distributed position within the simulation domain. Then, three outcomes are possible: i) the complex dissociates without nucleation, ii) a microtubule-bound nucleation, or iii) an unbound nucleation.

From the appearance point, *n* = 6 possible linear paths (or metatrajectories) are generated (Fig. 1E). We set *n* = 6 to reasonably consider the possible directions where nucleation complexes can diffuse. These paths are oriented in *n* equally spaced directions, with an overall offset that is chosen at random. Potential sites for microtubule-bound nucleation correspond to points where one of the paths intersects a microtubule at a distance *d*. The probability that a nucleation complex reaches a site at this distance from its appearance point without nucleating before is:

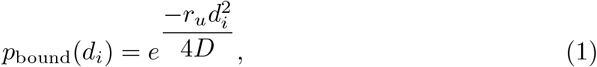

where *r*_*u*_ represents the rate of unbound nucleation, and *D* is the nucleation complex diffusion coefficient at the membrane. For computational efficiency, we only consider intersections at distances less than *d*_max_ = 1.5*µ*m, beyond which we consider *p*_bound_(*d*) to be zero based on our parameter choices. The probability of a potential microtubule-bound nucleation event is therefore given by the sum of *p*_bound_(*d*) over all directions where an intersection was found. Conversely, the probability of a potential isotropic nucleation event is

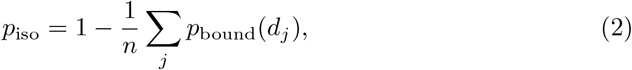

with the summation over *j* including only those paths that intersect a microtubule. We execute a single nucleation event either isotropic or on the first microtubule intersected by metatrajectory *i* based on these probabilities.

Based on empirical observation [40, 56], a complex reaching the lattice of a microtubule dissociates with probability 76%, while a freely diffusing complex dissociates with probability 98% (see [40]). Hence, before committing to either a microtubule-based or isotropic nucleation event, we reject the appearance event with a probability of 76% or 98%, respectively. These uniform post-hoc rejection probabilities are justified by the observation that membrane dissociation rates are very similar for free and microtubule-bound nucleation complexes that do not nucleate [40].

Isotropic nucleations are executed at the original appearance point. In case of a microtubule-based nucleation, a new microtubule is nucleated parallel, antiparallel, or branched to either side with an angular distribution with the mode at 35^*?*^ with respect to the parent microtubule, exactly as in [34], based on the experimental data by [41].

Previous observations showed that nucleation complexes move from gap regions to band regions as bands start emerging, with the total number of complexes remaining relatively constant [16]. To accommodate this observation, we model an overall fixed number of nucleation complexes *N*_*tot*_, which can be either free or occupied. Upon successful nucleation, a nucleation complex becomes occupied for a duration *t*_*occupied*_ = 60*s* (Fig. 1F), which is the average time until a nucleation complex is released from a new microtubule by katanin [56]. When handling an appearance event, the attempted nucleation is immediately rejected with a probability equal to the fraction of currently occupied complexes.

### 2.5 Protoxylem simulations

We modelled the local activity of proteins specifying the banded pattern (most likely ROPs and their downstream effectors [17, 19, 21]), using a difference in catastrophe rate between predefined band and gap regions as in [16]. Experimental observations of protoxylem development show that microtubule patterning starts from a well-established transversely oriented array [16, 49]. We, therefore, started our simulations with a two-part initiation phase similar to [16]. The first 30 minutes, all nucleations occurred at random positions with a variable angle *α*_*init*_, drawn from a normal distribution with a mean of *α*_*bias*_ and a variance of *α*_*noise*_ (Fig. 1H), to firmly establish array orientation. This was followed by 90 minutes of the nucleation mode for the remainder of the simulation, to generate a more realistic array microstructure. After the two hour initiation phase, we simulated protoxylem band formation by increasing the catastrophe rate in the gap regions by a factor *f*_*cat*_ (Fig. 1I), similar to simulations by Schneider et al. [16].

### 2.6 Expected mismatch angles

Although a ROP pattern is expected to follow the general orientation of the initial microtubule array, this match may not be exact, as the ROP pattern needs to wrap smoothly around the geometry, while maintaining an intrinsic band spacing [24]. We have previously shown [24] that the orientation of a spiral ROP pattern that maintains the distance between bands follows:

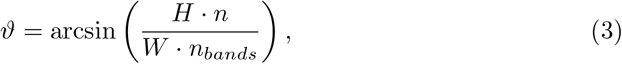

where *H* is the domain length, *W* the domain circumference, *n* the spiral number (1 for a single spiral, 2 for a double spiral, etc.), and *n*_*bands*_ is the number of bands in an equivalently spaced banded array. For our 10 bands, this equation gives a set of discreet angles that a ROP pattern is likely to follow. Assuming that the ROP pattern will always adopt the orientation closest to that of the microtubule array (reasonable at least for low spiral numbers; see [24]), microtubule arrays with orientations in between these discreet spiral angles will have the mismatch shown in Fig. 2E.

**Fig. 2.**
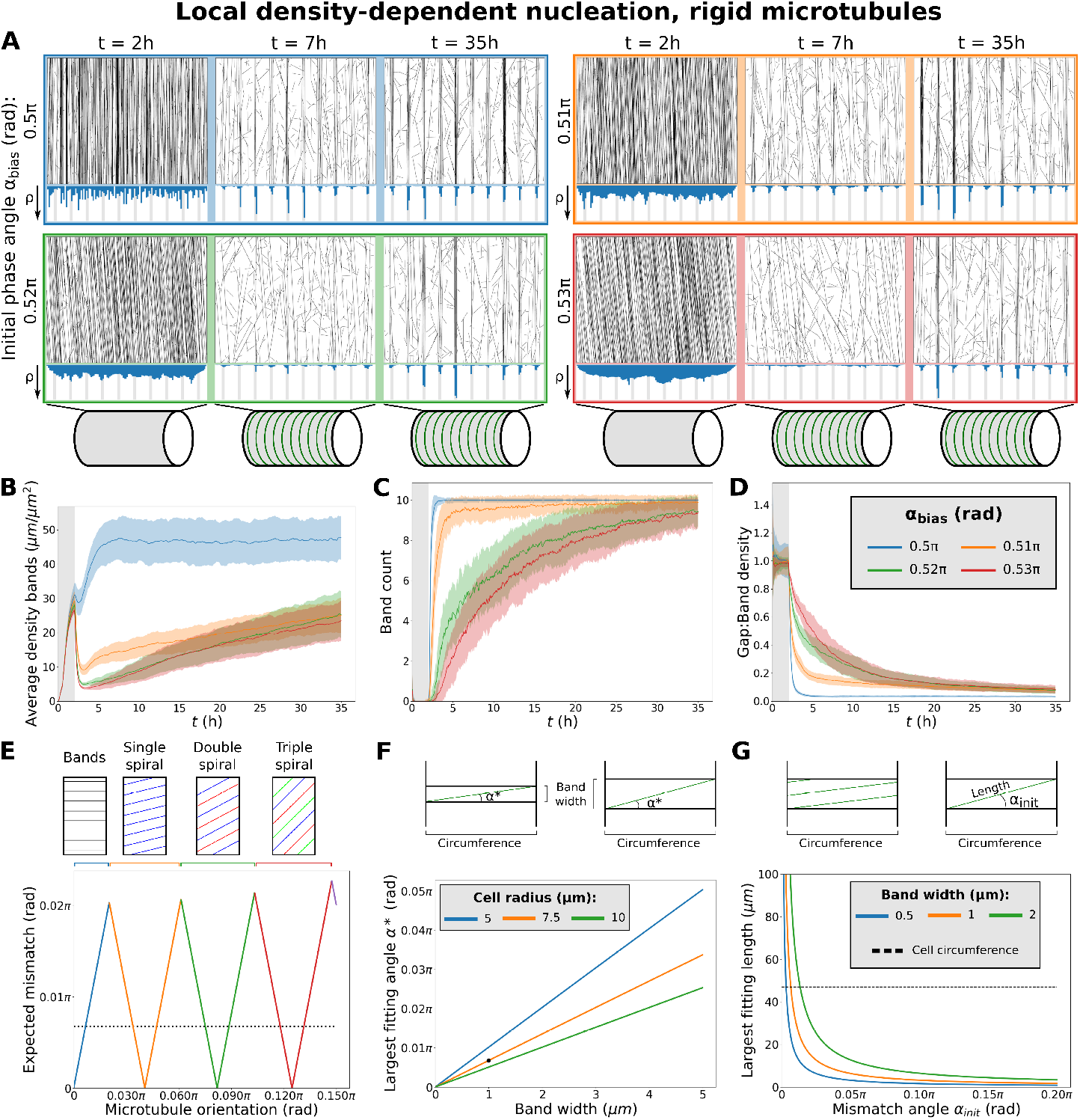
Fast protoxylem patterning is sensitively dependent on co-alignment between microtubules and the underlying pattern. (A) Snapshots from protoxylem simulations using starting arrays with bias angles *α*_*bias*_ of 0.5*π* (90°), 0.51*π* (91.8°), 0.52*π* (93.6°), and 0.53*π* (95.4°) in the first half hour with only minor deviations (*α*_*noise*_ = 0.01 *rad*^2^). Histograms below showing local microtubule density *ρ* share the same axis within a time series, but not among time series. (B) Average microtubule density in the band regions. (C) Number of populated bands, defined as bands with a microtubule density greater than three times the average density in the gaps. (D) Ratio of density in gaps and bands. Quantities in (B–D) were calculated from 100 simulations. The band formation phase starts at *t* = 2*h*, i.e., at the end of the grey area. Lines indicate the average and shaded areas the standard deviation. (E) Minimal expected mismatch angle of a spiral (or banded) ROP pattern following a microtubule array of different orientations based on geometrical constraints for a distance of 6 *µm* between the centres of bands and a cylindrical domain with a radius of 7.5 *µm*. Dotted line indicates the largest mismatch angle for which a microtubule can still span the cell’s circumference within a band. (F) Largest mismatch angle *α*^*∗*^ at which a microtubule (bundle) can fit within a band along the entire circumference of the cell. Black dot indicates default simulation values. (G) Largest length that a microtubule (bundle) can have while still fitting entirely within a band region at varying mismatch angles. Band width in simulations is 1 *µm* (orange line). Dashed line indicates cell circumference for comparison.

## 3 Results

### 3.1 Strong co-alignment of the microtubule array with a pre-existing band pattern facilitates rapid microtubule band formation

When starting with a microtubule array that is strongly co-aligned with the orientation of the band regions, we found that microtubule bands could form rapidly, both for isotropic nucleation (Fig. S.3 and S.4) and for local density-dependent nucleation (Fig. 2A–D). A slight mismatch of 3.6° (0.02*π* rad) in this co-alignment, however, already resulted in an extremely slow band formation process for both nucleation modes, similar to simulations in [16] with isotropic nucleations.

We observed that with a mismatch up to 1.8° (0.01*π* rad), the microtubule density in bands was maintained, whereas with mismatches of at least 3.6°, this density was lost at the beginning of the patterning process after which bands were “rediscovered” at a slow rate (Fig. 2B). These results suggest that timely microtubule band formation requires co-alignment between the initial microtubule array and the underlying pattern proposed to be formed by ROPs.

Some degree of co-alignment between the initial microtubule array and this underlying ROP pattern is biologically realistic, since the orientation of the microtubule array helps shape the orientation of the ROP pattern [19, 24]. However, the orientation of a ROP pattern is also influenced by geometrical constraints, as it has to form either rings or spirals that wrap around the circumference [24]. This requirement on the ROP pattern creates a minimal expected mismatch with the initial array, which, unlike the ROP pattern, is not limited to a discrete set of possible orientations. From this consideration, we calculated a minimal expected mismatch that can be as high as 3.6° (Fig. 2E). Our simulations suggested that this mismatch was too large for band formation without full breakdown and rediscovery (Fig. 2A–D).

As a proxy for calculating the maximum tolerable mismatch angle, we calculated the largest mismatch angle for which a straight microtubule could still span the circumference once while staying within the band (Fig. 2F) as well as the largest stretch of microtubule that could fit in a band given the mismatch angle (Fig. 2G). The point at which band formation began to suffer from the mismatch was similar to the point at which a single microtubule (bundle) could no longer stay in a band region along the entire circumference of the cell (Fig. 2F). For greater mismatch angles the largest length of microtubule (bundle) that could fit within a band without bending rapidly decreased (Fig. 2G). Variation of cell circumference and band width within the biologically relevant range suggests that the co-alignment requirement for straight microtubules is too strict to guarantee timely band formation.

The effect of co-alignment alone, therefore, did not explain timely band formation, and the use of more realistic microtubule nucleations did not improve upon this effect.

### 3.2 Microtubule flexibility can lead to density loss from bands under isotropic nucleation

Previous models with microtubule flexibility [37, 39] used values of *p* = 20–30 *µm* measured *in vivo* in animal cells [50, 51]. For microtubule dynamics parameters based largely on measurements in developing protoxylem [16], these persistence length values caused so many extra collisions that it resulted in a loss of density and alignment even in simulations without band formation (Fig. S.5A–C). Possibly, these persistence length values are too low for plant cells. Sasaki et al. [46] found average persistence lengths of 60 *µm* and 100 *µm* in their *in vitro* gliding assays with high and low microtubule densities, respectively. Measurements of microtubules suspended in flow cells even gave persistence lengths of around 2 *mm* for non-interacting microtubules [46]. Since the deflections due to microtubule interactions are modelled directly in our simulations, it is not just the ‘intrinsic’ persistence length that contributes to the measured persistence length and so the measured persistence length will be smaller than the value we should use to control deflections. Therefore, we investigated multiple larger persistence lengths, up to the millimetre range consistently measured in *in vitro* experiments with individual microtubules [45, 46]. For persistence lengths of hundreds of micrometers, aligned arrays did form (Fig. S.5A–C). Curiously, for these larger persistence lengths, the edge-induced catastrophes were not always sufficient to give the array a transverse orientation (Fig. S.5D). For our study of protoxylem band formation we, therefore, continued using the biased initiation phase.

Band formation was actually hindered by semiflexible microtubules when using isotropic nucleation. For *p* = 100 or 200 *µm*, stable starting arrays could be formed, but density was lost when band formation started (Fig. 3A and S.6). It would seem, therefore, that the extra flexibility, rather than helping microtubules find bands, actually makes microtubules already in bands bend out and suffer from the increased catastrophe rate in the gap regions. Proper bands only formed for more rigid microtubules with *p* = 500 or 1000 *µm*.

**Fig. 3.**
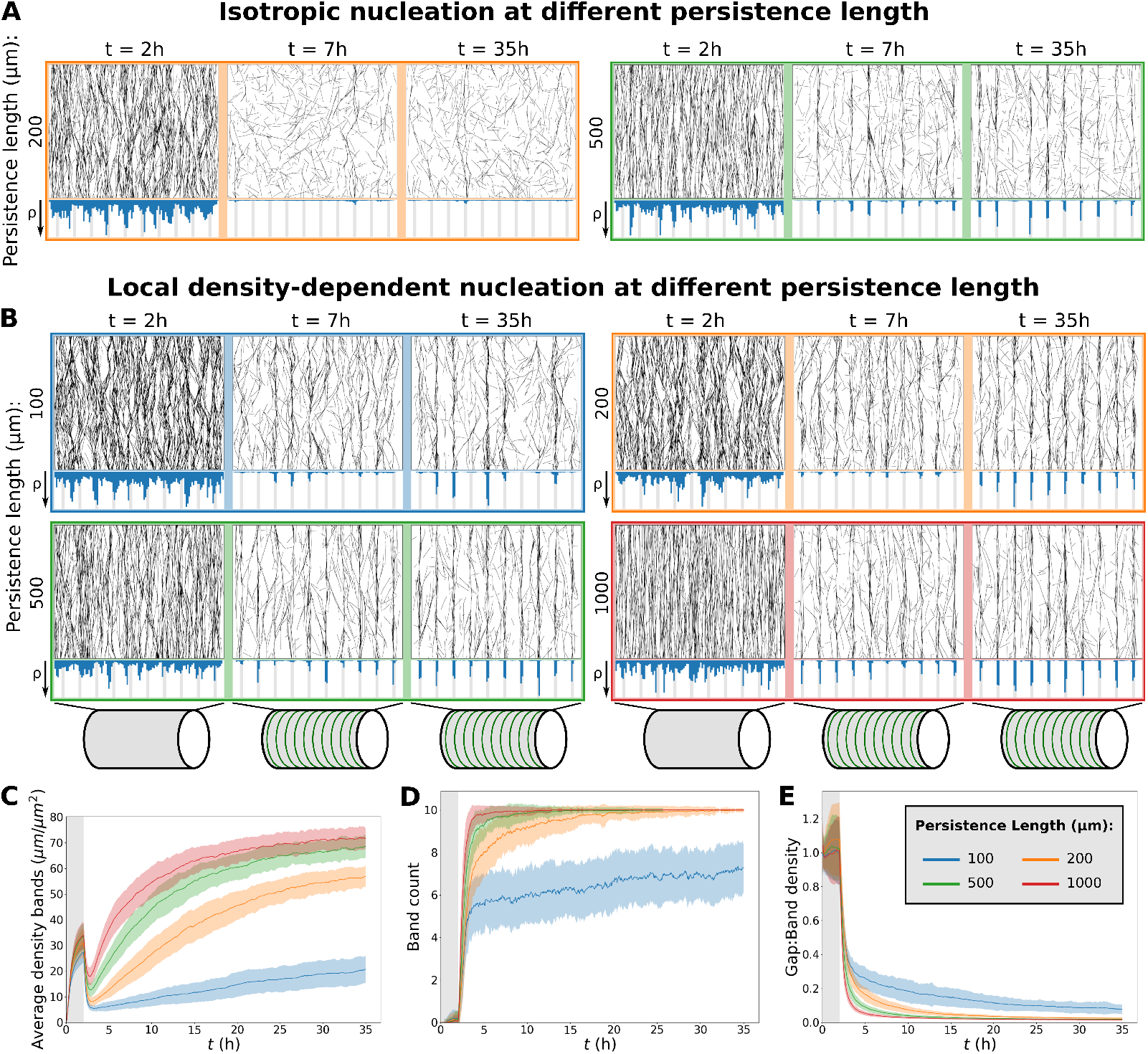
At lower persistence lengths too much density is lost to maintain bands. This problem is less severe local density-dependent nucleation than with isotropic nucleation. (A,B) Snapshots from protoxylem simulations with isotropic (A) and local density-dependent nucleation (B) for different microtubule persistence lengths. Starting arrays were obtained with transverse nucleations in the first half hour (*α*_*bias*_ = 0.5*π, α*_*noise*_ = 0.01 rad^2^). Histograms below showing local microtubule density *ρ* are plotted on the same scale within a time series, but not between time series. (C) Average microtubule density in the band regions. (D) Number of populated bands, defined as bands with a microtubule density greater than three times the average density in the gaps. (E) Ratio of density in gaps and bands. Quantities in (C–E) were calculated from 100 simulations with local density-dependent nucleation. The band formation phase starts at *t* = 2*h*, i.e., at the end of the grey area. Lines indicate the average and shaded areas the standard deviation.

The loss of density in the band regions at the lower persistence lengths could at least partially be counteracted when a significant portion of isotropic nucleations was moved to the gap regions (Fig. S.7). Such a shift could be expected to occur dynamically in cells as gap regions start to empty, when taking into account microtubule-bound nucleations, suggesting that a more realistic implementation of nucleations could be essential to band formation when taking microtubule flexibility into account.

### 3.3 Local density-dependent microtubule-based nucleation helps to keep bands populated with semiflexible microtubules, even for misaligned starting arrays

When combining semiflexible microtubules with local density-dependent nucleation, band formation improved for *p* = 100 *µm*, but most microtubule density was still initially lost, and took a long time to recover. However, band formation was now possible at *p* = 200 *µm*, lower than for isotropic nucleation, as nucleations were automatically allocated to the denser band regions as gaps density started to decrease (Fig. 3B–E). Furthermore, the combination of semiflexible microtubules and local density-dependent nucleation also greatly improved timely band formation for a significant mismatch between the orientations of the starting array and the band pattern. A mismatch as high as 18° (0.1*π* rad) in the angle of the nucleations in the initiation phase still yielded a partially banded pattern after five hours of band formation (Fig. 4A– D). As measured in Fig. 4F, our boundary conditions reduced the mismatch of the starting array compared to the bias angle. Still, the actual mismatch of corresponding to the 18° intended mismatch (*α*_*bias*_ = 0.6*π* rad) was about 10.8° (0.06*π* rad), substantially more than the worst mismatches we expected theoretically (Fig. 2E).

**Fig. 4.**
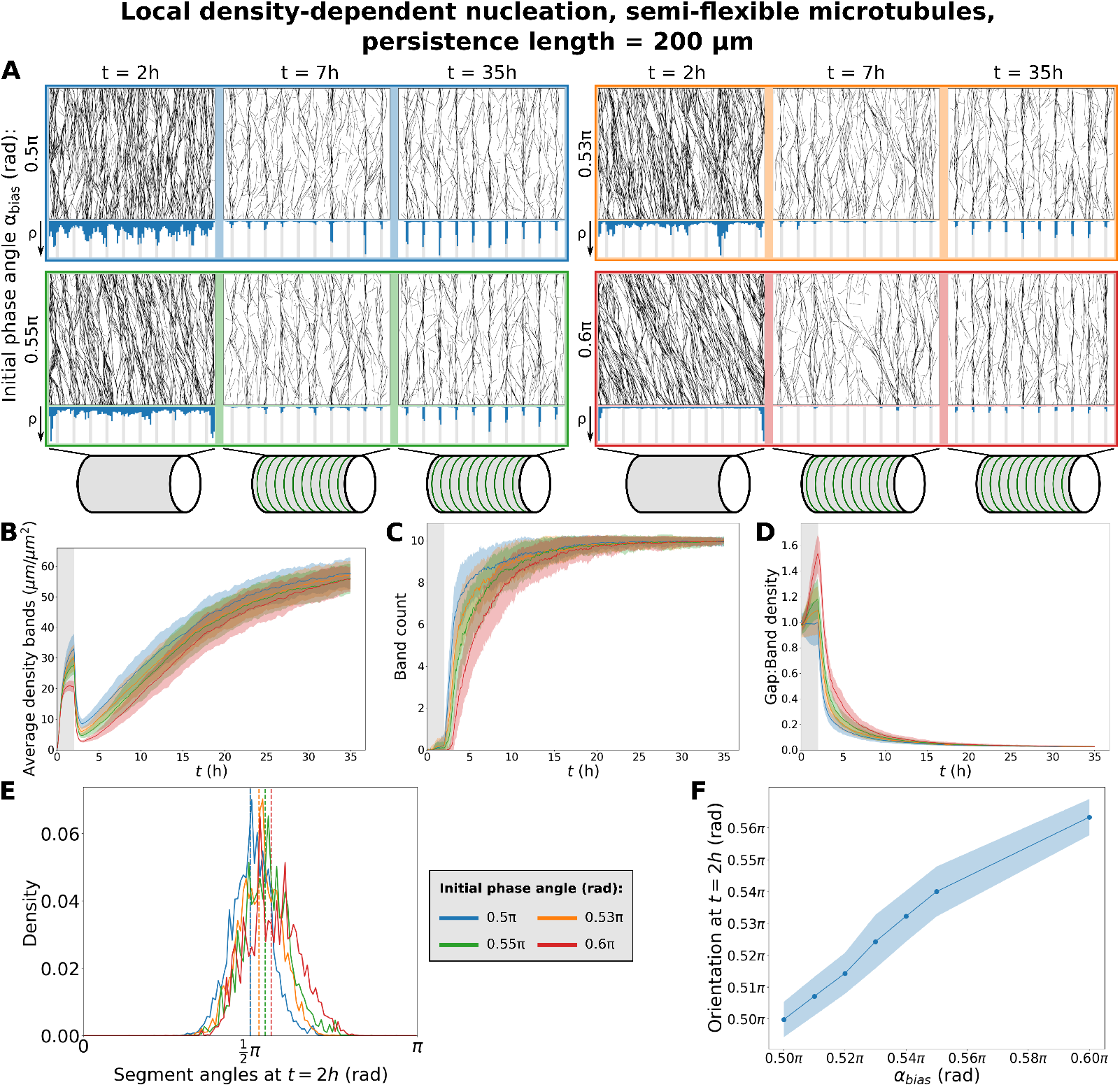
With local density-dependent nucleation (Fig. 1), greater mismatches in the initial orientation still allow fast band formation. (A) Snapshots from protoxylem simulations for *p* = 200 *µm* with local density-dependent nucleation using starting arrays with different bias angles *α*_*bias*_ in the first half hour with only minor deviations (*α*_*noise*_ = 0.01 rad^2^). Histograms below showing local microtubule density *ρ* are plotted on the same scale within a time series, but not between time series. (B) Average microtubule density in the band regions. (C) Number of populated bands, defined as bands with a microtubule density greater than three times the average density in the gaps. (D) Ratio of density in gaps and bands. (E) Distribution of microtubule segment angles, weighted by segment length, at *t* = 2*h* from the individual example simulations shown in (A). Dashed lines indicate the the overall array orientation. (F) Average array orientation at *t* = 2*h* as a function of the bias angle in the initiation phase. Quantities in (B–D), and (J) were calculated from 100 simulations. The band formation phase starts at *t* = 2*h*, i.e., at the end of the grey area. Lines indicate the average and shaded areas the standard deviation.

A reason for the increased tolerance for mismatches might be that the semiflexible microtubules inherently cover more different angles, of which a substantial portion could align with the band regions, even though the average orientation does not. This would give the array more opportunities to correct its course, by microtubules bending back into the array, or by ‘wrong’ orientations getting catastrophes in gap regions and rescues at a point where the orientation better matches the underlying ROP pattern. An investigation of the angles at the start of the band formation process in simulations with microtubule flexibility showed that there is indeed a broad distribution of microtubule segment angles (Fig. 4E).

## 4 Discussion

We have identified three important requirements for microtubule patterning in developing protoxylem by doing extensive simulations: (1) sufficient co-alignment between the microtubule array and the underlying ROP pattern, (2) moderate microtubule flexibility, and (3) realistic microtubule-based nucleation. Together, these aspects allow microtubule bands to form on a realistic time scale in fully interacting microtubule array simulations with idealised microtubule dynamics parameter values based on measured data [16].

One important aspect that we did not model explicitly is the interplay between microtubule patterning and ROP patterning. Here, we modelled the ROP pattern as a static, banded prepattern, whereas in reality, it is most likely shaped by a reaction diffusion process in which a combination of positive feedback activation and a difference in diffusion between active and inactive ROPs leads to spontaneous pattern formation (we extensively reviewed the details of such processes in plants elsewhere [57]). This ROP patterning has been modelled for both metaxylem [58] and protoxylem [24]. For protoxylem, the formation of the banded ROP pattern involves microtubules acting as ROP diffusion barriers that orient the pattern [24]. We partly modelled this orienting effect by generating starting arrays with various degrees of co-alignment. However, while real microtubule arrays generally align in a transverse orientation before the start of protoxylem patterning, they rarely have exactly the same orientation along the entire length [16, 49]. The resulting bands also tend to have a variation in their orientation and they are not all perfectly straight or equally spaced in ways that reflect initial densities and orientations in the starting arrays [16, 49]. These observations suggest that the underlying ROP pattern also provides co-alignment at a local level, which may help microtubule band formation, given moderate flexibility to follow the curved features. At the same time, the possibility of curved bands results in an increased rate of straight-growing microtubules leaving band regions, which we found could be detrimental to band formation. To help growing microtubules follow these curved bands, microtubule associated proteins (MAPs) involved in microtubule bundling may be important, e.g., MAP65-8, which is expressed in developing xylem [59]. In addition, microtubules must be sufficiently flexible to follow these curves.

Our simulations demonstrated the importance of microtubule flexibility even when assuming straight band and gap regions. Flexibility improved the ability of microtubules to follow predefined band regions in spite of small mismatches in the orientation of the microtubules and the band regions. However, our simulations also indicated a trade-off, where increased flexibility means more microtubules curve into gap regions where they are more likely to suffer catastrophes. Microtubule associated proteins may, however, reduce this effect in practice by preventing microtubules from bending into the gap regions.

The need for microtubule flexibility becomes even more obvious when we consider metaxylem patterning, where arrays need to form circular or ellipsoid gaps. In microscopic pictures, microtubules appear to curve around these gaps [60, 61]. The gapped structure also means that microtubule patterning cannot rely on co-alignment with the ROP pattern as in protoxylem patterning. Therefore, metaxylem patterning may require additional proteins to help the microtubules form this structure that are absent or less important in protoxylem patterning. Recent evidence indicated the involvement of microtubule-associated protein MAP70-5, which lowers microtubule persistence lengths, enabling the formation of microtubule loops [46, 61]. This is consistent with earlier observations that MAP70-5 lines the borders of the pits and its overexpression increases the ratio of pitted to spiral cell wall patterns [62]. CORTICAL MICROTUBULE DISORDERING (CORD) proteins may also be involved. These proteins disorder microtubules by partially detaching them from the membrane [60], possibly facilitating corrections in the microtubule orientation. Other potential factors include ‘Boundary of ROP domain 1’ (BDR1), Wallin, and actin networks that form in the pits, though these may only act in a later stage, during formation of the pit borders [63].

In addition to the importance of microtubule flexibility, our simulations also showed the importance of realistic microtubule-based nucleation. We have previously shown the importance of locally saturating microtubule-based nucleation for array homogeneity in Jacobs et al. [40]. There, we assumed a constant, uniform supply of nucleation complexes as an important source of local saturation. Here, we relaxed this assumption by allowing microtubules to draw from a global pool of nucleation complexes, while maintaining a local density-dependence of the nucleation rate. Consequently, the reduction of microtubule density in the gap regions would free up nucleation complexes that could then boost the nucleation rate in the bands. This effect helped compensate for microtubule loss from catastrophes suffered by microtubules leaving the band region. This partial shift in the location of nucleation complexes is in line with microscopic observations of nucleation complexes during protoxylem development [16] and upon oryzalin treatment [40].

Another interesting finding was that the introduction of a finite persistence length made it harder to obtain global alignment from edge-induced catastrophes alone. This reduction of global alignment compared to local alignment has been observed before with rigid microtubules on large domains [64], but microtubule flexibility reduced the domain size at which locally aligned patches formed. Microtubule flexibility, therefore, increases the importance of local orienting cues, such as the observed sensitivity of microtubules to mechanical stress [65–72]. It has also been suggested that, depending on the spacing of microtubule-membrane linkers, individual microtubules may also tend towards a longitudinal orientation to minimize their bending energy before anchoring [73, 74]. It remains to be seen, however, whether this mechanical force would be strong enough to overrule the orienting forces arising from collective microtubule interactions [42].

The lowest persistence lengths reported in Sasaki et al. [46], *≈* 30*µm* in high density assays including MAP70-5, are comparable to the 20-30 *µm* measured in animal cells [50, 51]. The largest persistence length in gliding assays in Sasaki et al. [46], *≈* 100*µm* in low density assays without MAP70-5, was on the low side for timely band formation, but there remains a large gap with the millimetre range values they observed in flow cells. This difference suggests a large uncertainty in the value of the microtubule persistence length in plant cells, as well as a possibility for cells to modulate it. In absence of sufficient membrane attachment, strong microtubule bending can be induced by forces generated by active cellulose synthase complexes [75] and cytoplasmic streaming [54, 76]. Therefore, good candidates for persistence length modulation are proteins involved in microtubule-membrane linkage, such as CELLULOSE-MICROTUBULE UNCOUPLING (CMU) proteins [75], and certain IQ67-Domain (IQD) proteins [77], of which IQD13 functions in metaxylem development [23]. IQD proteins can also be regulated dynamically, in particular through calcium signalling, as they have calmodulin-binding domains [78, 79]. This kind of regulation may dynamically influence microtubule persistence length. Therefore, persistence lengths measured in one situation may not necessarily apply to the next, making it necessary to obtain *in vivo* persistence length measurements for plant cells of different types.

## 5 Conclusion

In conclusion, we have shown that co-alignment, microtubule flexibility, and local density-dependent nucleation are important aspects of protoxylem patterning. Our findings lay the groundwork for future studies on patterns generated by microtubule-ROP interactions. These studies may include simulations of other systems, such as metaxylem, as well as simulations that combine existing ROP models (e.g., [24]) with microtubule simulations.

## 6 Author contributions

B.J., M.S., J.M., L.F. and E.E.D. designed the research. B.J., M.S. and E.E.D. performed the research. B.J. analysed data. B.J., M.S. and E.E.D. wrote the paper. B.J., M.S., J.M., L.F. and E.E.D. reviewed and edited the paper.

## 9 Data and Coding Availability

We are in the process of producing a new release of CorticalSim, which will be made public along with the publication of [42]. The release will become accessible via [80].

## Appendix A Simulation details and parameter values

Simulations were performed with an extended version of ‘CorticalSim’ [80], a well established and fast two-dimensional microtubule simulation platform [16, 31, 34, 36, 44, 52]. To account for the possibility that some katanin severing events at crossover intersections may have been interpreted as induced catastrophes [36] in the original experiments and analysis by Dixit and Cyr [28], the probability *P*_*cat*_ that collisions at large angles result in catastrophes was lowered compared to previous studies [31, 34]. Contrary to those studies, we included katanin severing by default, with a rate of *r*_*x*_ = 0.023 *s*^*−*1^ per crossover, similar to values found in experiments [29, 48, 81].

Protoxylem simulations were performed with 1 *µm* wide band regions separated by 5 *µm* wide gap regions as in Schneider et al. [16]. Simulations started with a 2*h* initiation phase without bands followed by a 5*h* or 33*h* band formation phase in which the catastrophe rate in the gap regions was increased by a factor *f*_*cat*_ = 3, which in experimental observations tends to be achieved or exceeded during a substantial part of the patterning process [16]. In the first 0.5*h* of the initiation phase, nucleations were distributed uniformly and given an angle *α*_*bias*_, with a small amount of normally distributed noise, with a variance *α*_*noise*_ of 0.01 rad^2^. For simulations with rigid microtubules, this biased phase had almost no collisions that could lead to induced catastrophes or crossovers. To compensate, the nucleation rate for this phase was reduced by a factor 4 for simulations with rigid microtubules. The remainder of the initiation phase used the same nucleation mode as the band formation phase. Simulations without bands were run for 7*h* starting from an empty array with additional isotropic nucleations added at the beginning speed up the population of the array. These *‘seeded’ nucleations* were added at a density of 0.1 *µm*^*−*2^*s*^*−*1^ and a rate of 0.003 *s*^*−*1^ as in Lindeboom et al. [43]. See Table A1 for default parameter values.

**Table A1.**
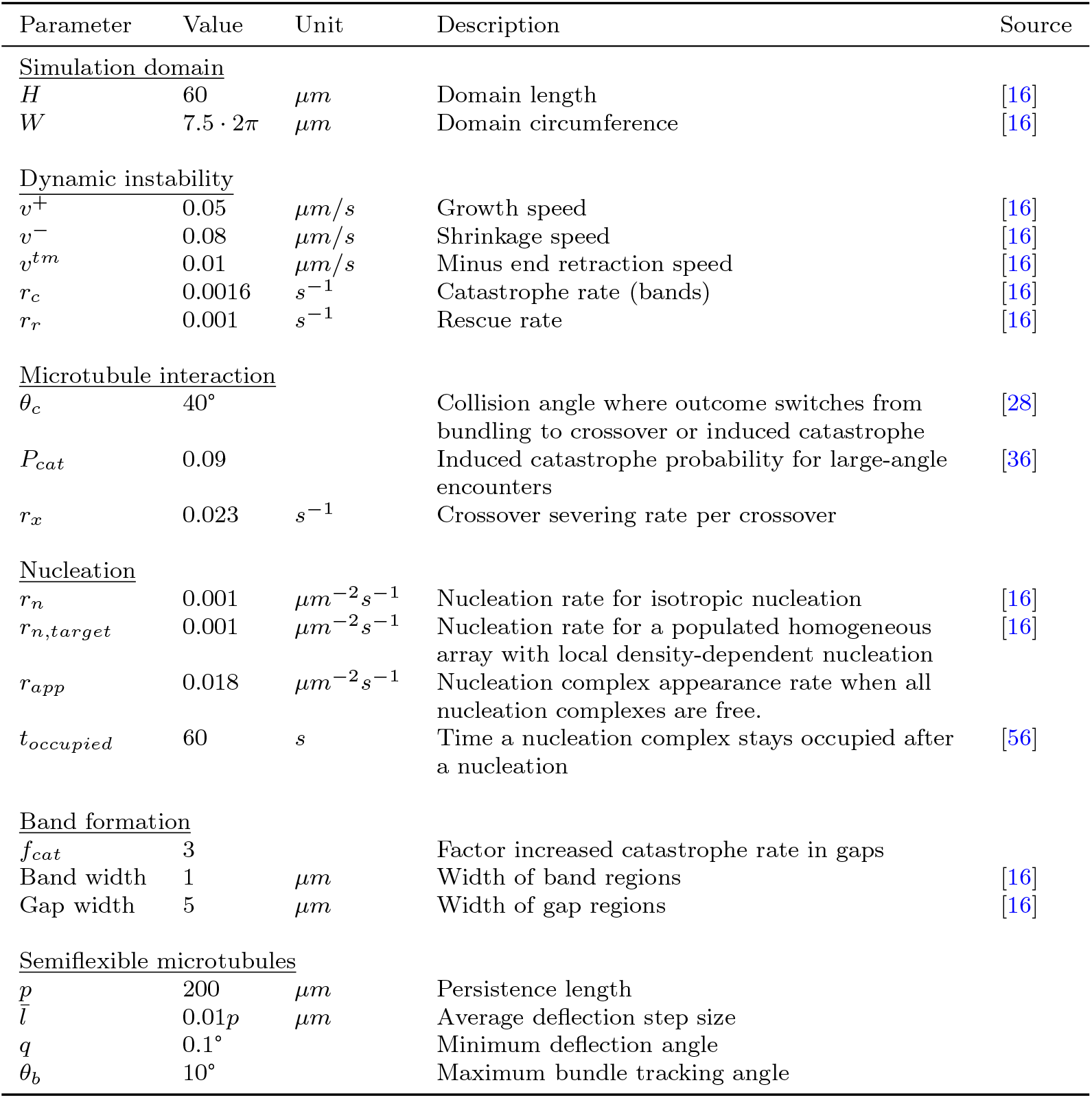
Default parameter values used in the simulations.

### A.1 Target nucleation rate

For calculating nucleation parameters, we aimed at an overall target nucleation rate *r*_*n,target*_ for homogeneous arrays of 0.001 nucleations *µm*^*−*2^*s*^*−*1^, consistent with previous work [16, 34, 40]. As a first estimate, we assumed that the fraction of unbound nucleations in a fully populated array would be negligible.

From there, we calculated back to a parameter *r*_*n,max*_ (the nucleation rate when all nucleation complexes are free and all nucleations microtubule-bound). For this, we used the factor difference *f*_*rn*_ between the nucleation rate when all complexes are available and the target nucleation rate. We estimated this rate from the ratio between microtubule-associated appearances in nearly empty oryzalin-treated arrays (0.013 appearances *µm*^*−*2^*s*^*−*1^) and total, mostly microtubule-associated, appearances in established arrays (0.0037 appearances *µm*^*−*2^*s*^*−*1^) that were measured in [40]. This approach gives:

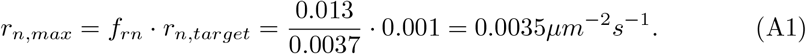

With a rejection probability of 0.76 for microtubule-bound nucleations, we computed a required maximum appearance rate of:

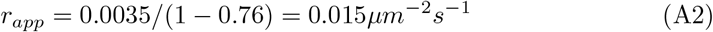

We further used a fixed duration *t*_*occupied*_ of 60s during which a nucleation complex remains occupied upon nucleation, based on an average of 58.9s observed in experimental work [56]. Using our estimates for parameters *r*_*n,target*_, *r*_*n,max*_, and *t*_*occupied*_, we calculated the remaining parameter *N*_*tot*_. At nucleation rate *r*_*n,target*_, a number of *N*_*occ,target*_ nucleation complexes are expected to be occupied, following:

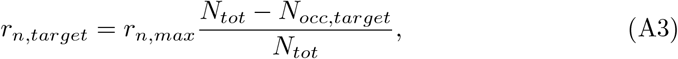

where *N*_*tot*_ is the total number of nucleation complexes. Using this expression and the fact that the number of occupied complexes depends on the duration of occupancy *t*_*occupied*_ and the rate at which they become occupied (i.e., the global nucleation rate), we found the following expression for *N*_*tot*_:

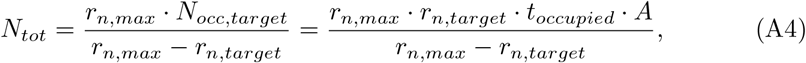

where *A* is the total domain area.

From initial simulations of homogeneous arrays, we found that this approach resulted in a realised nucleation rate of about 80% of *r*_*n,target*_ (Fig. S.1). To keep the realised nucleation rate close to the target, we increased the appearance rate by 20% to 0.018 *µm*^*−*2^*s*^*−*1^.

## Appendix B Semiflexible microtubules

To implement semiflexible microtubules, we adapted an approach from Mirabet et al. [37]. We gave microtubules deflections in their growth direction at discrete points, separated by variable distances *l* drawn from an exponential distribution with mean 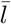 . Deflection angles were drawn from a uniform interval [*−m, m*], where *m* was computed to obtain the desired persistence length given 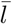 (see next section). Angles of which the absolute value was smaller than the minimum deflection angle *q* = 0.1° were set to zero to avoid numerical problems. For 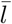 we chose a value of 1% of the desired persistence length *p* by default. This value prevents large numbers of very small deflections from resulting in many very similar trajectories that would needlessly slow down simulations and may give technical difficulties, while still keeping the step size small relative to the persistence length.

For individual microtubules averaged over sufficient length, the length of the average deflection step size 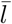 does not matter for the persistence length *p* of the total microtubule as long as the appropriate deflection angle is used. However, microtubules in a populated array interact with each other, and so there may well be a difference between many small deflections and fewer larger deflections. Therefore, we tested the effect of different step sizes for the same persistence length, for a range of step sizes feasible in the current simulation setup (Fig. S.2). For the same persistence length, the deflection step size seemed to have little effect on the alignment and orientation of arrays without bands (Fig. S.2D,E). For lower persistence lengths in particular, there seems to be a small effect on the overall density, likely resulting from differences in the rates of encounters that could lead to crossover-severing [36] or induced catastrophes (Fig. S.2A–C). Therefore, the precise way in which microtubules are flexible, may also have some impact on the array as a whole, but the magnitude of this impact on array alignment, orientation, and density is limited.

## Appendix C Persistence length calculations

Persistence length *p* measures how fast the correlation between the orientation of two different points on a microtubule decays with the microtubule length between these points. We use the following definition of persistence length *p*:

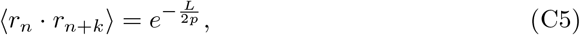

where ⟨*r*_*n*_ *r*_*n*+*k*_⟩is the average inner product of unit vectors *r*_*i*_ in the direction of the microtubule at points *n* and *n* + *k* and *L* is the length along the microtubule between these two points.

Since Eq. C5 holds for any two points *n* and *n* + *k*, and we are using independent deflections, it is sufficient to look at a single deflection after length *l* between points *n* and *n* + 1:

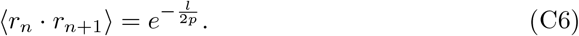

Without loss of generality, we assume the arbitrary initial angle (of *r*_*n*_) to be 0. For a given deflection angle *ϑ* (Fig. C1A) *r*_*n*+1_ then is given by:

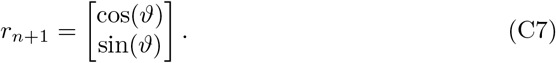

**Fig. C1.**
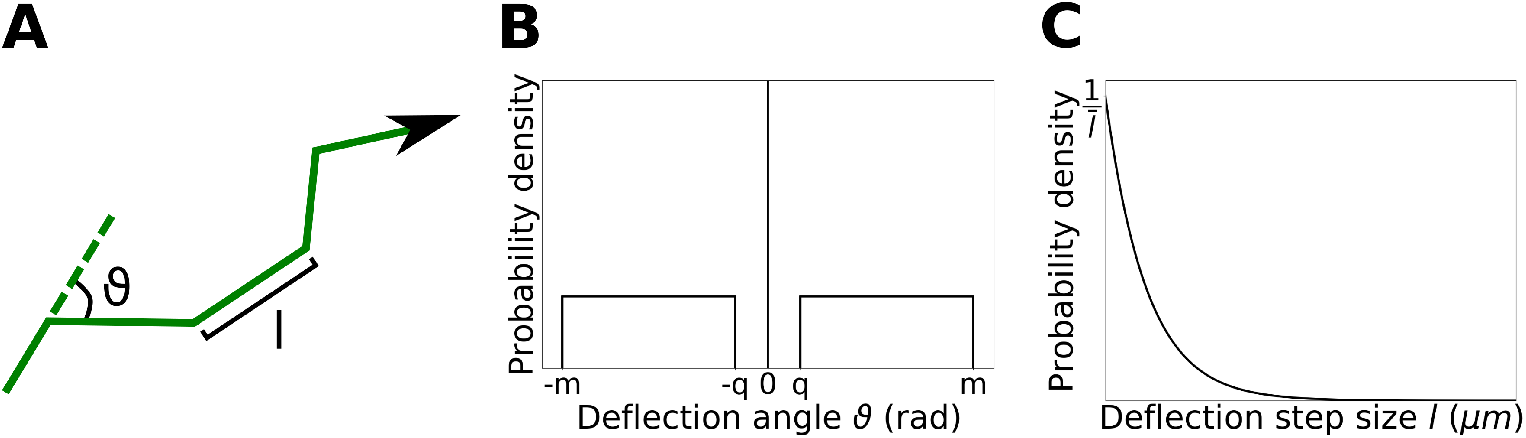
Semiflexible microtubule implementation details. (A) Simulated microtubules get deflections of angle *ϑ* every deflection step size *l*. Angles in cartoon have been exaggerated for visibility. (B) Deflection angles are drawn from a uniform interval [*−m, m*], with all angles in [*−q, q*] set to zero. Value of *q* in the graph is exaggerated for visibility. (C) Deflection step sizes are drawn from an exponential distribution with average ^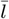^ .

For the inner product in Eq. C6, we then get:

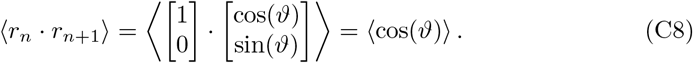

We take deflection angles *ϑ*, drawn from a uniform interval [*−m, m*], with the smallest angles (|*ϑ*| *< q*) set to zero for numerical reasons (Fig. C1B). In that case, we get:

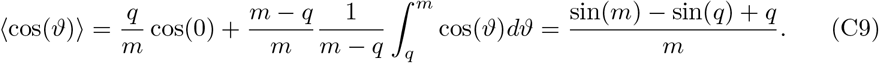

Taking deflection step lengths *l* drawn from an exponential distribution with mean 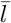 (Fig. C1C), we can now determine the persistence length *p* for a given value of *m*:

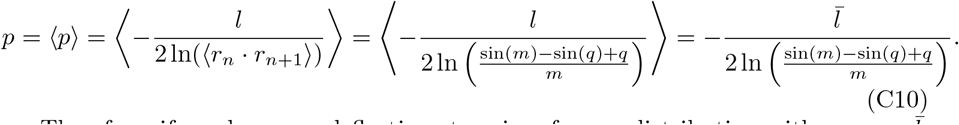

Therefore, if we draw our deflection step sizes from a distribution with average 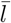, we can solve the boundary value *m* of the interval from which we draw the deflection angles from the above equation for the desired persistence length *p*. This means that we can control the microtubule persistence length in our simulations using 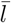, *m*, and *q* as input parameters, with one of *m* and 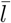 calculated to obtain the desired *p*.

## Appendix D Summary statistics

Array alignment was quantified using the planar nematic order parameter *S*_2_ as commonly used in polymer physics and often used for quantifying cortical microtubule alignment [31, 34, 52]:

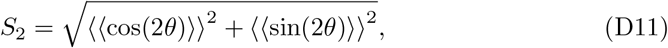

where *θ* is the angle of individual microtubule segments with the *x*-axis of the simulation domain, and double angular brackets indicate a length-weighted average over all microtubule segments. Alignment parameter *S*_2_ is zero for a completely isotropic array and one for a perfectly aligned array (ignoring microtubule polarity).

Overall array orientation Θ was also computed as commonly used in simulation studies [34, 52]:

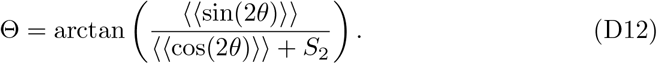

Note that both these quantities are computed on the unrolled cylinder mantle of the simulation domain, i.e., as if it were a flat, strictly two-dimensional sheet.

Band counts were calculated as the number of bands with a microtubule density at least three times higher than the average density in the gaps. This measure was selected after testing multiple options for best representing visual inspections of array snapshots and histograms.

Averages and standard deviations of orientations from multiple simulation runs were calculated using the average and standard deviation for angular quantities. The average orientation 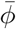 over *N* simulations is:

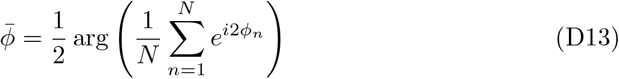

where arg is the argument of a complex number, and *ϕ*_*n*_ is the orientation of the *n*^*th*^ simulation. The factor 2 in the exponent and factor 1/2 before the argument correct the fact that we consider orientations without direction (so in [0, *π*) rather than [0, 2*π*)). The circular standard deviation *sd*_*ϕ*_ is:

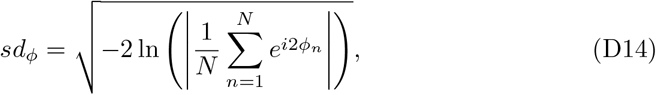

where ln is the natural logarithm and the vertical bars indicate the absolute value of a complex number.

