## Supplementary figures for "Protoxylem microtubule patterning requires ROP pattern co-alignment, realistic microtubule-based nucleation, and moderate microtubule flexibility"

### Supplementary information for: Microtubule flexibility enhances the patterning potential of the plant cortical microtubule array

Bas Jacobs, Marco Saltini, Laura Filion, Jaap Molenaar, Eva E. Deiunum

#### 1 Supplementary figures

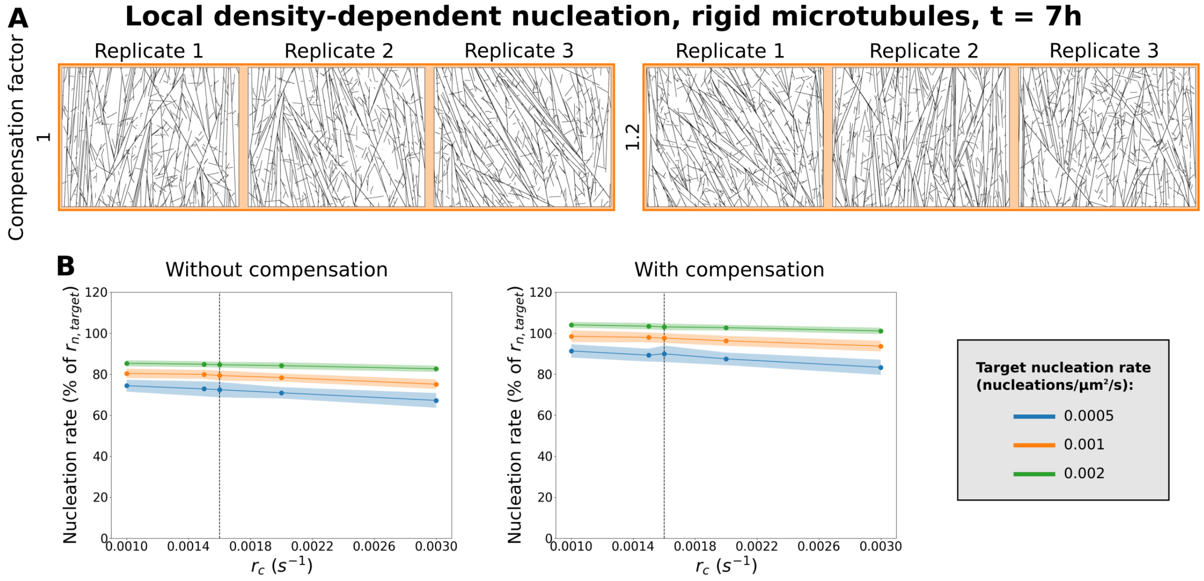


**Figure S.1: A twenty percent compensation on the nucleation complex appearance rate keeps the achieved nucleation rate close to the target.** (A) Snapshots at $t = 7h$ from array simulations without bands with locally saturating nucleation with and without compensation. (B) Realised nucleation rate, calculated over a 200 s measurement interval, as a percentage of the target rate ($r_{n,target}$). Quantities in (B) were calculated from 100 simulations. Lines indicate the average and shaded areas the standard deviation.

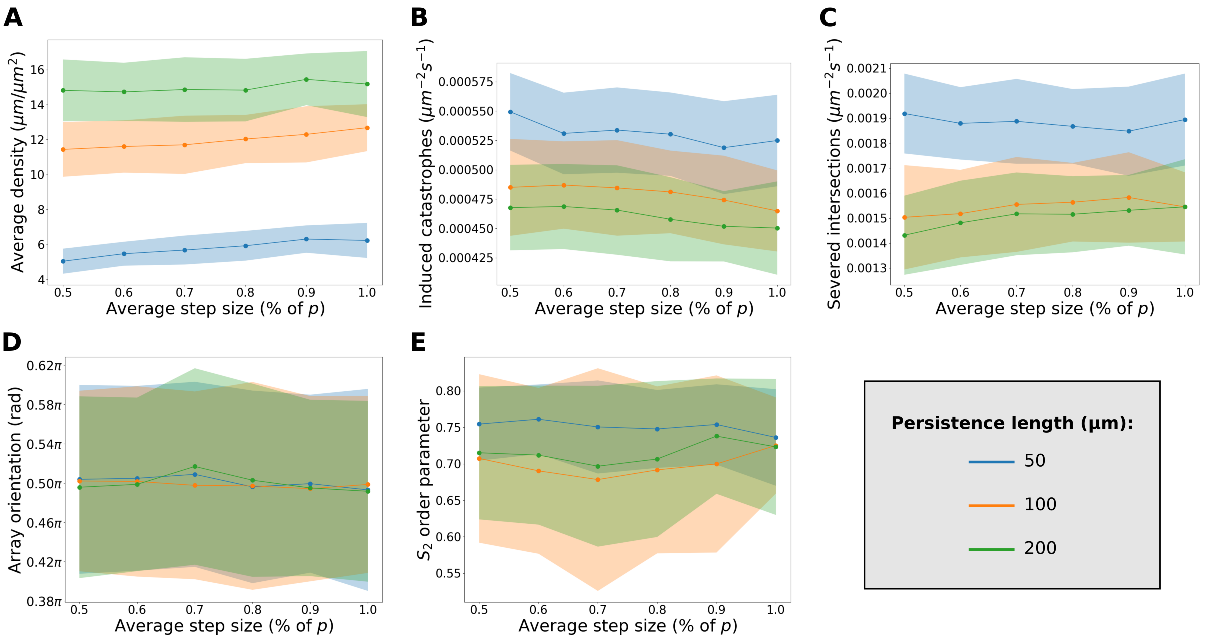


**Figure S.2: Influence on array behaviour of different step sizes for the same persistence length.** (A–E) Quantification of array state and simulation processes at $t = 7h$ from simulations without bands using isotropic nucleation for various difference persistence lengths and deflection step sizes. Simulations used the default cylindrical geometry and were initiated with seeded nucleations as described in Appendix A. (A) Average microtubule density. (B) Overall number of induced catastrophes per unit area per second. (C) Overall number of intersection severing events per unit area per second. (D) Average array orientation. (E) $S_2$ order parameter, showing degree of alignment. (A,D,E) Quantities measured at $t = 7h$ intervals. (B,C) Quantities averaged over the last 200 s measurement interval. Quantities in (A–E) were calculated from 100 simulations. Lines indicate the average and shaded areas the standard deviation. Step sizes ($\bar{l}$) are expressed as a percentage of the persistence length $p$.

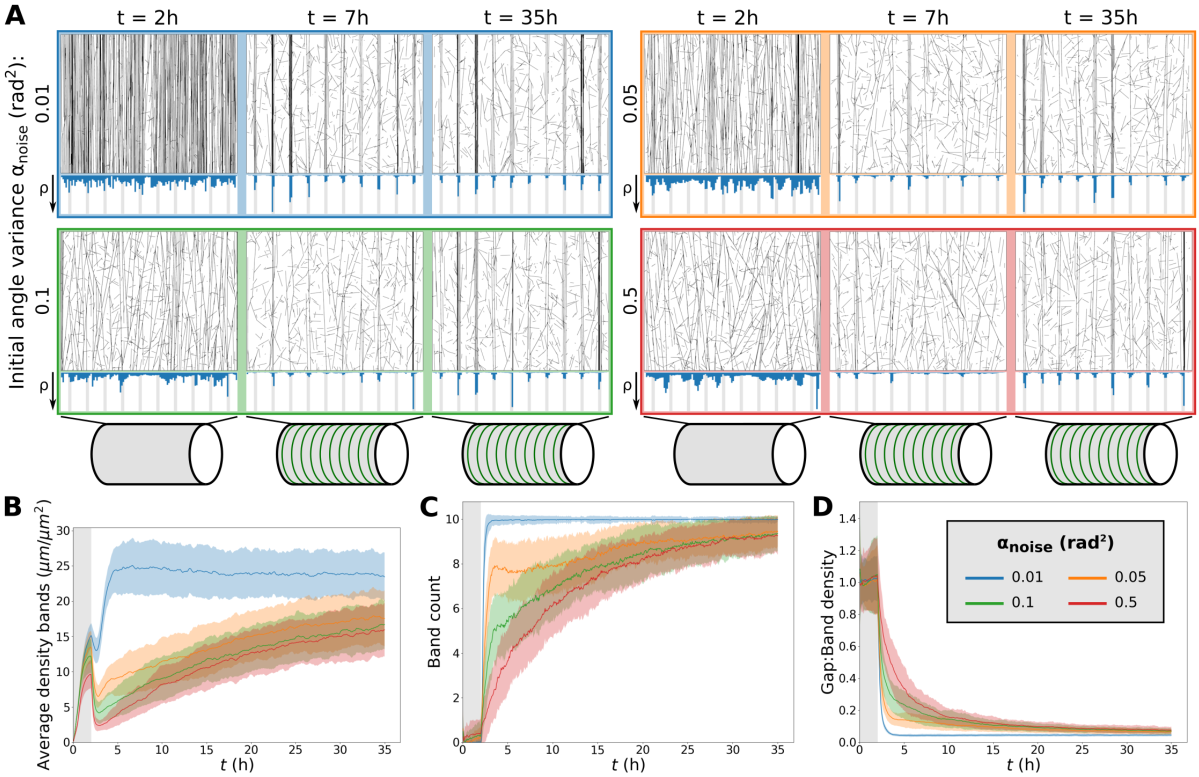


**Figure S.3: Isotropic nucleation allows fast band formation for sufficiently co-aligned arrays.** (A) Snapshots from protoxylem simulations with isotropic nucleation using starting arrays obtained with different values for $\alpha_{noise}$. Histograms below snapshots showing local microtubule density $\rho$ share the same axis within a time series, but not among time series. (B) Average microtubule density in the band regions. (C) Number of populated bands, defined as bands with a microtubule density greater than three times the average density in the gaps. (D) Ratio of density in gaps and bands. Quantities in (B–D) were calculated from 100 simulations. The band formation phase starts at $t = 2h$, i.e., at the end of the grey area. Lines indicate the average and shaded areas the standard deviation.

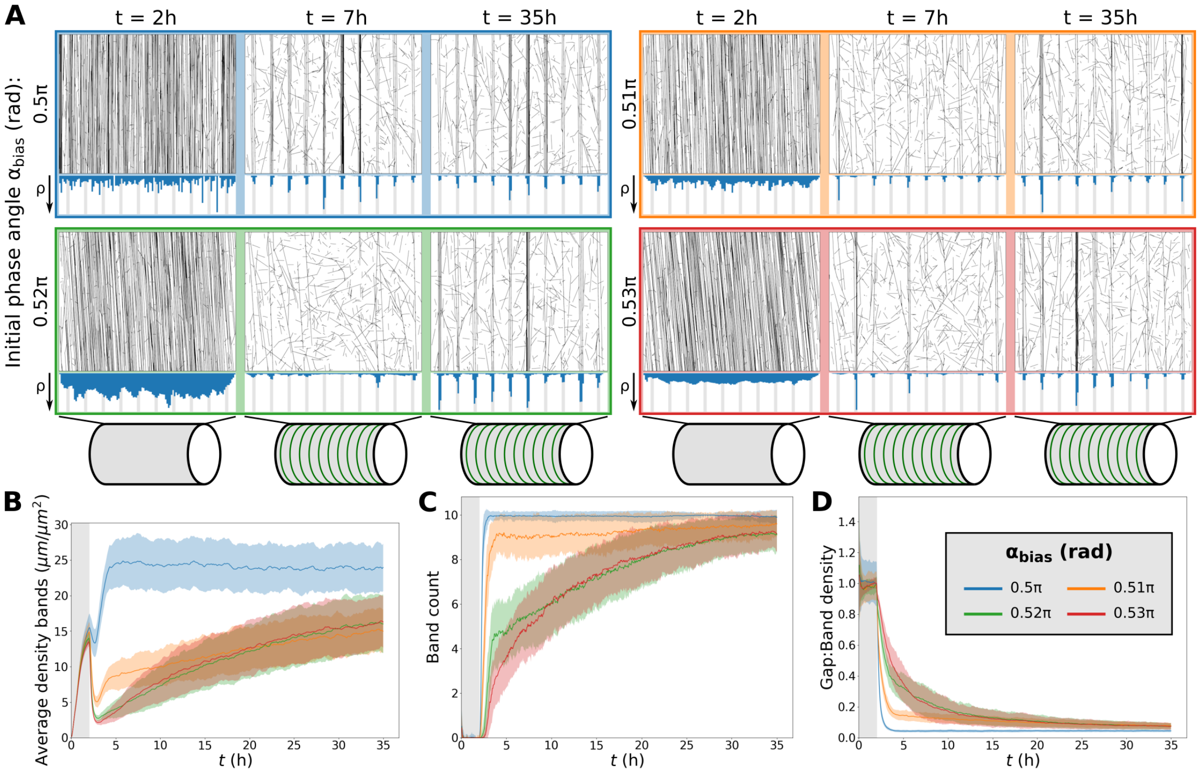


**Figure S.4: Fast protoxylem patterning is sensitively dependent on co-alignment between microtubules and the underlying pattern.** (A) Snapshots from protoxylem simulations with isotropic nucleation using starting arrays with different bias angles $\alpha_{bias}$ in the first half hour with only minor deviations ($\alpha_{noise} = 0.01\ rad^2$). Histograms below showing local microtubule density $\rho$ share the same axis within a time series, but not among time series. (B) Average microtubule density in the band regions. (C) Number of populated bands, defined as bands with a microtubule density greater than three times the average density in the gaps. (D) Ratio of density in gaps and bands. Quantities in (B–D) were calculated from 100 simulations. The band formation phase starts at $t = 2h$, i.e., at the end of the grey area. Lines indicate the average and shaded areas the standard deviation.

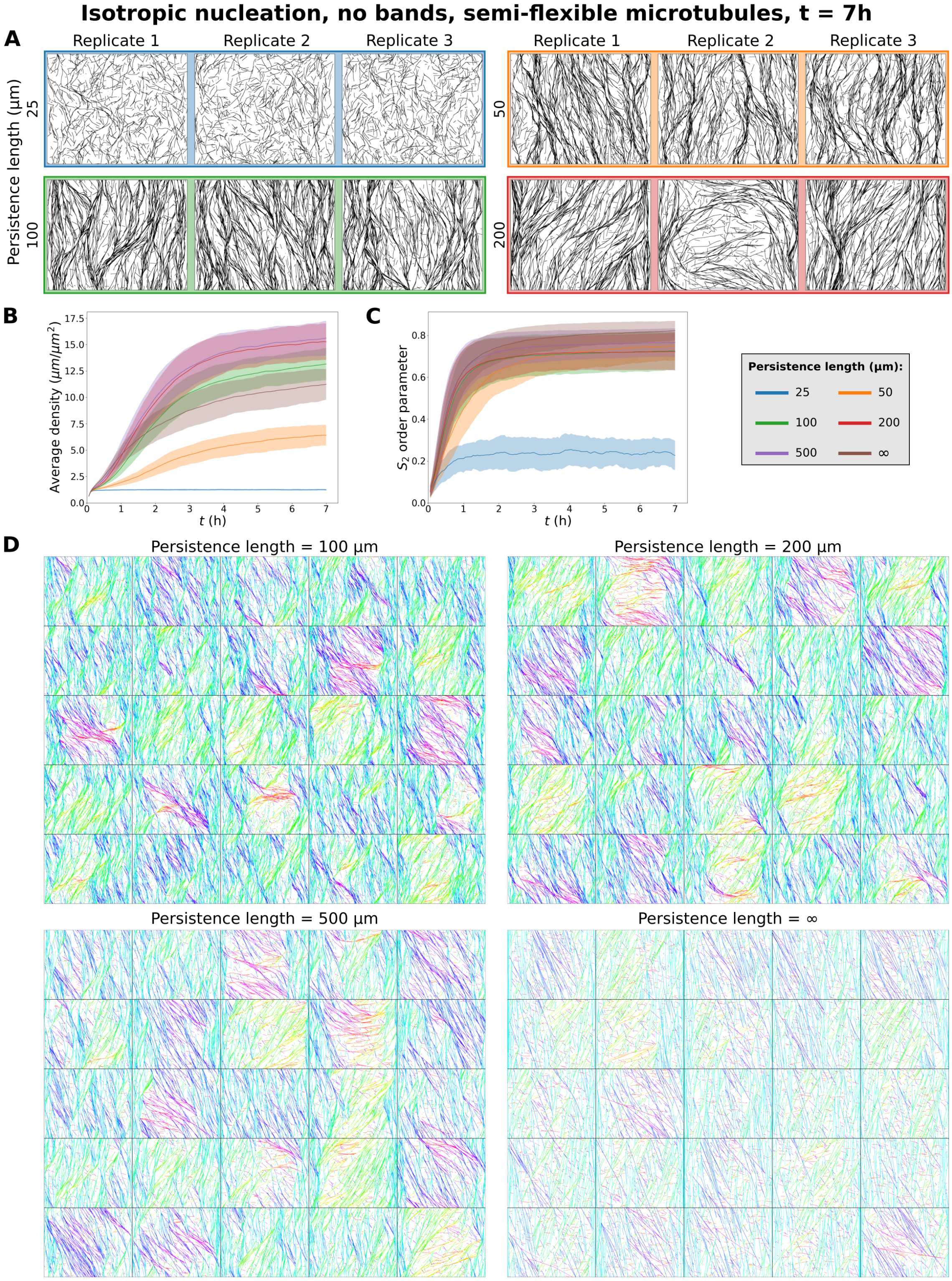


**Figure S.5: Semiflexible microtubules result in low densities and failure to align at low persistence lengths and reduced global alignment at intermediate persistence length.** (A) Snapshots at $t = 7h$ from array simulations without bands with isotropic nucleation for different microtubule persistence lengths. (B) Average microtubule density. (C) $S_2$ order parameter, showing degree of alignment. Quantities in (B–C) were calculated from 100 simulations. Lines indicate the average and shaded areas the standard deviation. (D) Snapshots at $t = 7h$ from 25 independent array simulations with isotropic nucleation without bands for different persistence lengths, and rigid microtubules (infinite persistence length). Microtubule segments are coloured by their orientation.

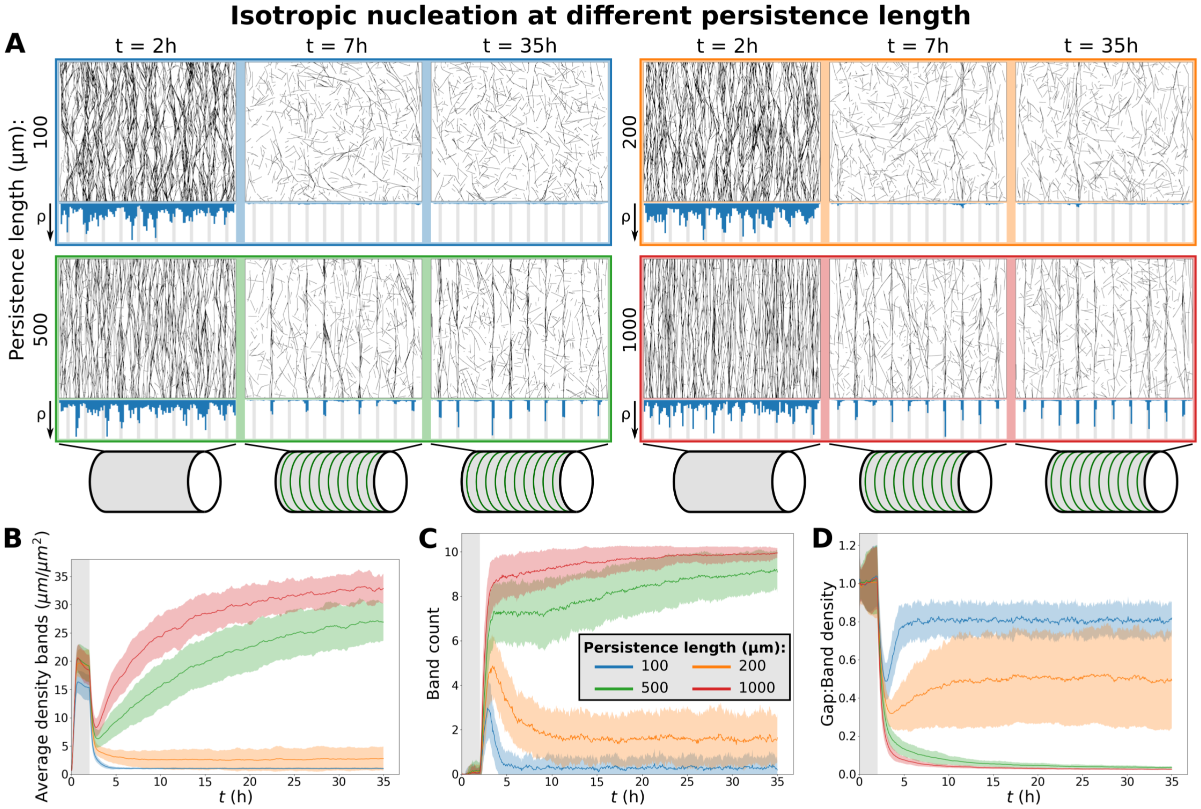


**Figure S.6: Band maintenance under isotropic nucleation requires high persistence length.** (A) Snapshots from protoxylem simulations with isotropic nucleation for different microtubule persistence lengths. Histograms below showing local microtubule density $\rho$ are plotted on the same scale within a time series, but not among time series. Starting arrays were obtained with transverse nucleations in the first half hour ($\alpha_{bias} = 0.5\pi$, $\alpha_{noise} = 0.01$ rad$^2$). (B) Average microtubule density in the band regions. (C) Number of populated bands, defined as bands with a microtubule density greater than three times the average density in the gaps. (D) Ratio of density in gaps and bands. Quantities in (B–D) were calculated from 100 simulations. The band formation phase starts at $t = 2h$, i.e., at the end of the grey area. Lines indicate the average and shaded areas the standard deviation.

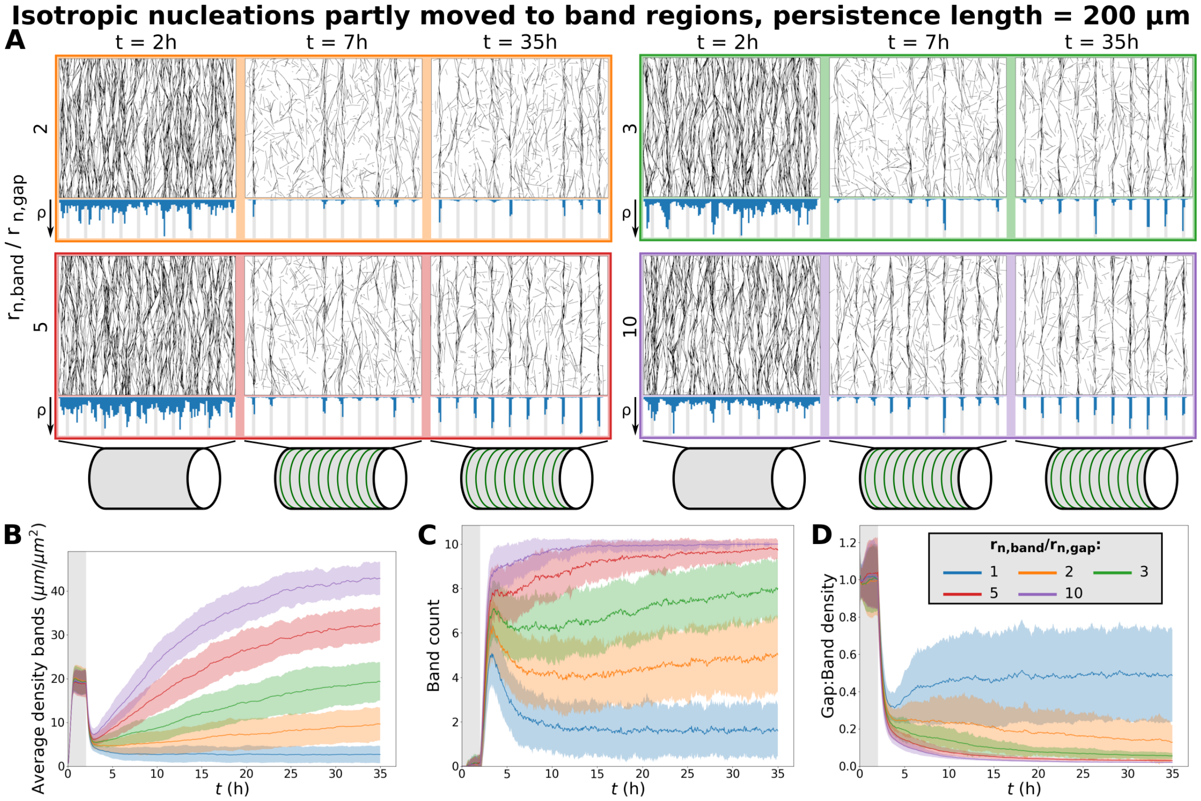


**Figure S.7: With isotropic nucleation, a shift of nucleations from gaps to bands can maintain band density.** (A) Snapshots from protoxylem simulations with a fraction of isotropic nucleations moved from gaps to bands, keeping the overall nucleation rate the same. The persistence length used was 200 $\mu m$. Starting arrays were obtained with transverse nucleations in the first half hour ($\alpha_{bias} = 0.5\pi$, $\alpha_{noise} = 0.01\ rad^2$). Histograms below showing local microtubule density $\rho$ are plotted on the same scale within a time series, but not among time series. (B) Average microtubule density in the band regions. (C) Number of populated bands, defined as bands with a microtubule density greater than three times the average density in the gaps. (D) Ratio of density in gaps and bands. Quantities in (B–D) were calculated from 100 simulations. The band formation phase starts at $t = 2h$, i.e., at the end of the grey area. Lines indicate the average and shaded areas the standard deviation.
